# Task-evoked pupil responses reflect internal belief states

**DOI:** 10.1101/275776

**Authors:** O. Colizoli, J.W. de Gee, A.E. Urai, T.H. Donner

## Abstract

Perceptual decisions about the state of the environment are often made in the face of uncertain evidence. Internal uncertainty signals are considered important regulators of learning and decision-making. A growing body of work has implicated the brain’s arousal systems in uncertainty signaling. Here, we found that two specific computational variables, postulated by recent theoretical work, evoke boosts of arousal at different times during a perceptual decision: decision confidence (the observer’s internally estimated probability that a choice was correct given the evidence) before feedback, and prediction errors (deviations from expected reward) after feedback. We monitored pupil diameter, a peripheral marker of central arousal state, while subjects performed a challenging perceptual choice task with a delayed monetary reward. We quantified evoked pupil responses during decision formation and after reward-linked feedback. During both intervals, decision difficulty and accuracy had interacting effects on pupil responses. Pupil responses negatively scaled with decision confidence prior to feedback and scaled with uncertainty-dependent prediction errors after feedback. This pattern of pupil responses during both intervals was in line with a model using the observer’s graded belief about choice accuracy to anticipate rewards and compute prediction errors. We conclude that pupil-linked arousal systems are modulated by internal belief states.

## Introduction

Many decisions are made in the face of uncertainty about the state of the environment. A body of evidence indicates that decision-makers use internal uncertainty signals for adjusting choice behavior^1–3^, and deviations between expected and experienced rewards for learning^4,5^. The brain might utilize its arousal systems to broadcast such computational variables to circuits implementing inference and action selection^4,6–9^.

Recent theoretical work postulates two variables at different moments during a challenging perceptual decision^1,10,11^: (i) decision confidence before feedback (i.e., the internally estimated probability of a choice being correct, given the available evidence) and (ii) prediction error (i.e., the difference between expected and experienced reward) after receiving feedback. Critically, and different from previous work on reinforcement learning^7,8,12^, the prediction error signals depend on graded internal confidence^10^ rather than on the categorical stimulus identity (see *Model Predictions* below). Such internal belief states have been incorporated in more recent models of reinforcement learning^13^. Building on previous work implicating arousal in uncertainty monitoring^3,14–16^, we here asked whether these two computational variables would evoke responses of central arousal systems.

It has long been known that the pupil dilates systematically during the performance of cognitive tasks, a phenomenon referred to as task-evoked pupil response^17–24^. Physiological work indicates that non-luminance mediated changes in pupil diameter are closely linked to central arousal state^25–28^. We quantified pupil responses during a perceptual choice task combined with reward-linked feedback, analogous to the task used in recent monkey work on uncertainty and prediction errors^10^. We then compared pupil responses before and after reward-linked feedback to predictions derived from alternative computational models of the internal variables encoded in the brain. Our goal was to (i) replicate the previously found scaling of pupil responses with decision uncertainty before feedback^3^ and (ii) test for the same scaling of pupil responses after feedback, as observed for dopamine neurons^10^.

## Results

We monitored pupil diameter in 15 human participants performing an up vs. down random dot motion discrimination task, followed by delayed reward-linked feedback (Figure 1). The random dot motion task has been widely used in the neurophysiology of perceptual decision-making^29,30^. Importantly, our version of the task entailed long and variable delays between decision formation and feedback, enabling us to obtain independent estimates of the pupil responses evoked by both of these events. We titrated the difficulty of the decision (by varying the evidence strength, or motion coherence, see Methods), so that observers performed at 70% correct in 2/3 of the trials in one condition (‘Hard’) and at 85% correct in 1/3 of the trials in the other condition (‘Easy’). Correct vs. error feedback was presented after choice and converted into monetary reward, based on the average performance level across a block (25 trials), as follows: 100% correct yielded 10 Euros, 75% yielded 5 Euros, chance level (50% correct) yielded 0 Euros. The total reward earned (in Euros) was presented on the screen to participants at the end of each block.

**Figure 1.**
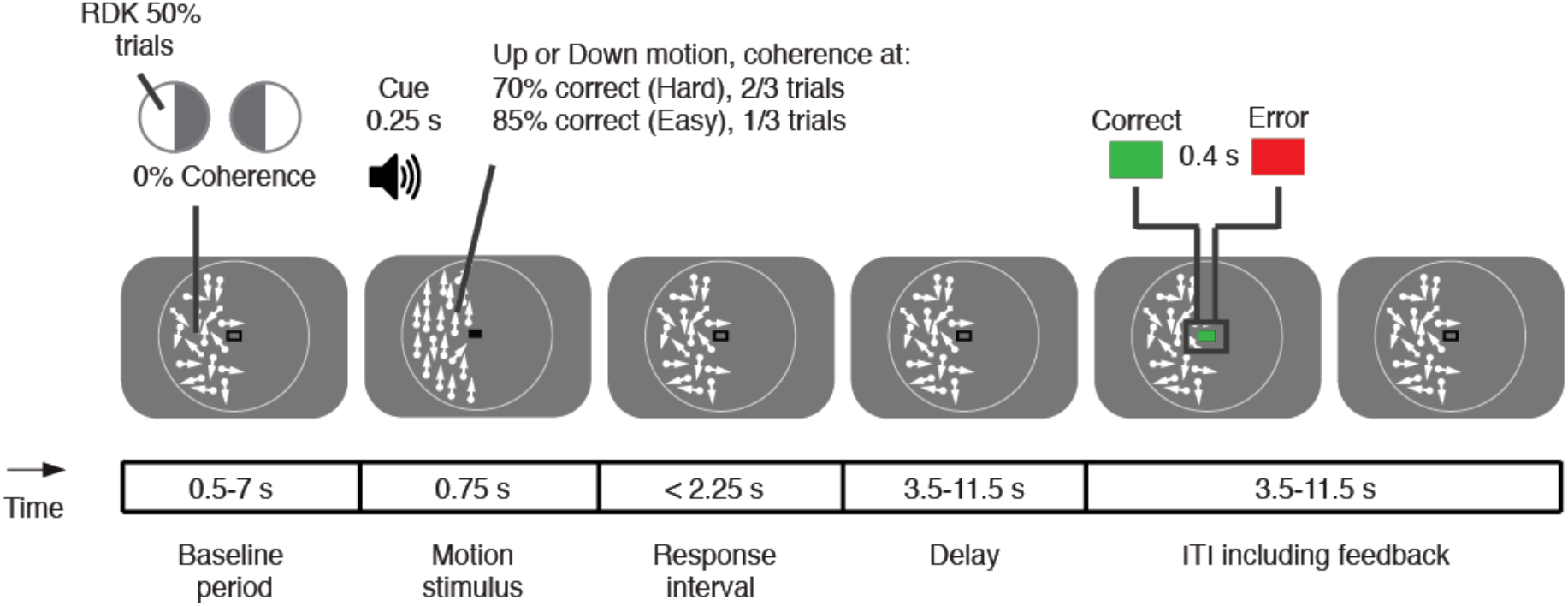
Perceptual choice task with delayed reward. Random dot kinematograms (RDK) were presented in one half of the visual field during each block of trials (counterbalanced). Random motion (0% coherence) was presented throughout all intervals except for the ‘motion stimulus’ interval, during which the RDKs to be discriminated were shown, prompted by an auditory cue (250 ms). Motion coherence of the stimulus varied from trial to trial, yielding a Hard and an Easy condition. A change from an open to a closed rectangle in the fixation region (constant luminance) prompted subjects’ choice (‘response interval’). After a variable delay (3.5-11.5 s) following the choice, feedback was presented that was coupled to a monetary reward (see main text). The white circle surrounding the RDKs is for illustration only and was not present during the experiment.

### Model predictions

We used two computational models based on signal detection theory^31^ to generate qualitative predictions for the behavior of internal signals before and after reward feedback that might drive pupil-linked arousal (Figure 2a, see Materials and Methods for details). Both models assumed that observers categorize the motion direction based on a noisy decision variable, which in turn depended on the stimulus strength (Hard or Easy), the stimulus identity (Up or Down), and on internal noise. The models’ choices were governed by comparing this noisy decision variable to zero, ensuring no bias towards one over the other choice.

**Figure 2.**
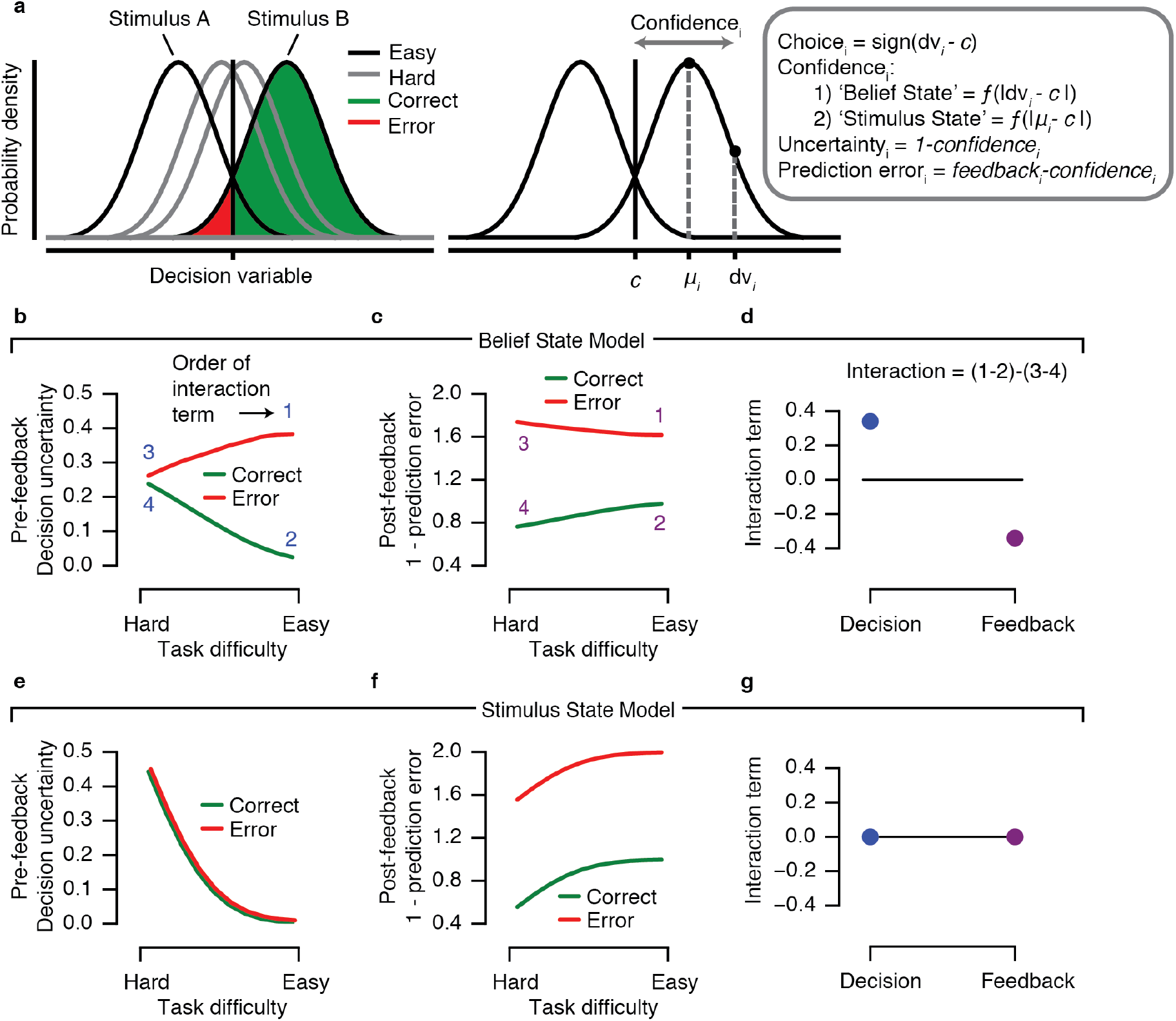
Alternative predictions for internal signals during pre- and post-feedback intervals of the task. **(a)** Computations underlying choice, confidence, uncertainty and prediction error. Repeated presentations of a generative stimulus produce a normal distribution of internal decision variables (*dv*) due to the presence of internal noise, which is centered around the generative stimulus (*μ*). In this model, confidence is defined as the single-trial distance between *dv* and *c*, the internal decision bound. Prediction errors are computed by comparing experienced reward (i.e. feedback) with the observers’ expected outcome. **(b-d)** Computational variables were simulated for every trial, then averaged separately for Correct and Error conditions for each level of task difficulty (in this case, motion coherence). Belief State Model predictions. **(e-g)** Stimulus State Model predictions. **(b, e)** Decision uncertainty (complement of confidence) as function of task difficulty during pre-feedback interval. **(c, f)** Prediction error as function of task difficulty during post-feedback interval. **(d, g)** Interaction term computed as (Easy Error - Easy Correct) - (Hard Error - Hard Correct). See main text for model details.

The two models differed in how confidence was defined. Here, with confidence we refer to the observer’s internally estimated probability that a choice was correct given the available evidence^11^. Because choice accuracy was coupled to a fixed monetary reward in our experiment (see above), confidence equaled an ideal observer’s internally estimated probability of obtaining the reward, in other words, reward expectation. In the ‘Belief State Model’, confidence was computed as the absolute distance between the decision variable (depending on the stimulus identity, stimulus strength, and internal noise) and the decision criterion (i.e., zero) (Figure 2a; see Methods and ref.^1^). By contrast, in the ‘Stimulus State Model’, confidence was computed as the absolute distance between the physical stimulus value (i.e. physical stimulus identity times stimulus strength) and the criterion (zero). In both models, reward prediction error was computed as the difference between the confidence and the reward-linked feedback. Thus, in the Belief State Model, the observer’s internal belief about the state of the outside world (encoded in the noisy decision variable) determined both reward expectation (i.e., confidence) and reward prediction error; in the Stimulus State Model, these computational variables did not depend on the observer’s internal belief, but only on the strength and identity of the external stimulus.

We simulated these two models to derive qualitative predictions that distinguished between their internal signals. To this end, we computed confidence and reward prediction errors at the level of individual trials (see above) and then collapsed these single-trial signals within each Accuracy and Difficulty condition. The rationale was that the interaction between conditions (defined as [Easy Error - Easy Correct] - [Hard Error - Hard Correct]) most clearly dissociated between the predictions generated from both models (Figure 2b-g).

Previous pupillometry work on a similar task showed that pre-feedback pupil responses scaled with decision uncertainty (i.e. the complement of decision confidence)^3^. We thus generated predictions for decision uncertainty during the pre-feedback interval (Figure 2b,e) and, by analogy, for the complement of prediction error during the post-feedback interval (Figure 2 c,f).

The critical observation is that the Belief State Model predicts a positive Accuracy x Difficulty interaction pre-feedback, and a negative interaction post-feedback (Figure 2d). This pattern is consistent with predictions from a reinforcement learning model based on a partially observable Markov decision process (POMDP)^10^. In contrast, the Stimulus State Model does not predict an Accuracy x Difficulty interaction either pre- or post-feedback (Figure 2g). This pattern is consistent with traditional reinforcement learning models^7,8,12^.

Previous work on perceptual choice has shown that reaction time (RT) scales with decision uncertainty^3,32,33^, in line with the Belief State Model. The same was evident in the present data: There was a main effect of accuracy, *F*_(1,14)_ = 51.57, *p* < 0.001, and difficulty, *F*_(1,14)_ = 19.53, *p* < 0.001, as well as an interaction effect of both, *F*_(1,14)_ = 34.95, *p* < 0.001, on RT (see Supplementary Fig. S1, compare with Figure 2b), in line with the Belief State Model. This indicates that, in our current data, a graded, noisy decision variable similar to the one postulated by the Belief State Model was encoded and used for the decision process. We next tested which of the two models better reflected responses of pupil-linked arousal systems. We analyzed pupil responses as a function of motion coherence and choice correctness for the two critical intervals of the trial: the phase of reward anticipation before feedback, as in previous work^3^, and critically, the phase of reward prediction error signaling after feedback.

### Sustained pupil response modulations during pre- and post-feedback intervals

The pupil responded in a sustained fashion during both intervals: after the onset of the motion stimulus and locked to the observers’ reported choice (i.e., pre-feedback) and post-feedback (Figure 3a, blue and purple lines). The pupil response remained elevated during feedback anticipation, long after stimulus processing (maximum of 3 s, 0.75 s stimulus duration plus response deadline of 2.25 s, see Figure 1). Upon feedback presentation, the pupil initially constricted due to the presentation of the visual feedback stimulus (see Supplementary Fig. S2) and then dilated again to a sustained level for the remainder of the post-feedback interval. Please note that we subtracted the pupil diameter during the pre-feedback period from the feedback-locked responses (see Methods), so as to specifically quantify the feedback-evoked response.

**Figure 3.**
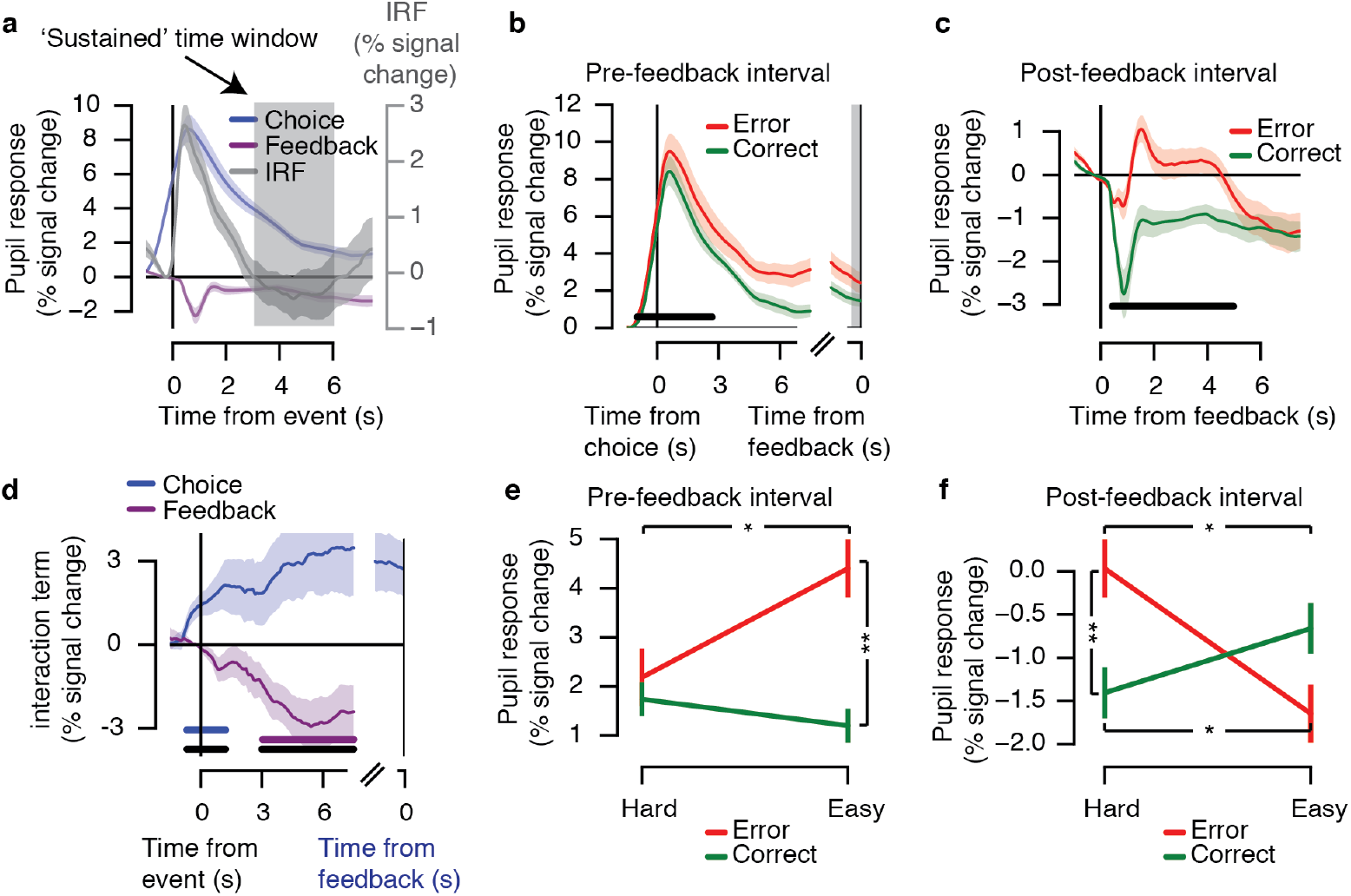
Pupil responses before and after feedback reflect observers’ belief state. **(a)** Pupil responses locked to the observer’s reported choice (blue) and locked to feedback (purple). Shown for comparison is the pupil ‘impulse response’ from same participants (IRF, see main text). Grey shading indicates sustained time window (3-6 s), in which IRF returned to baseline. **(b)** Evoked pupil responses for Correct and Error trials in pre-feedback interval. Black bar indicates correct vs. error effect, *p* < 0.05 (cluster-based permutation test). **(c)** Evoked pupil responses for Correct and Error trials in post-feedback interval. Black bar indicates correct vs. error effect, *p* < 0.05 (cluster-based permutation test). **(d)** Interaction term (Easy Error - Easy Correct) - (Hard Error - Hard Correct) for choice-locked (blue, coinciding with observers’ reported choice) and feedback-locked (purple) responses. Horizontal bars indicate effect, *p* < 0.05 (cluster-based permutation test): blue bar indicates choice-locked response tested against 0; purple bar indicates feedback-locked response tested against 0; black bar indicates difference in interaction between both responses. **(e)** Mean response in the pre-feedback interval (500 ms preceding feedback), as a function of difficulty and accuracy. **(f)** Mean response (in sustained time window) during the post-feedback interval, as function of difficulty and accuracy. Error bars, standard error of the mean (*N* = 15). \**p* < 0.05, \*\**p* < 0.01, ***p < 0.001.

For comparison, we measured, in the same participants (separate experimental blocks), pupil responses evoked during a simple auditory detection task (button press to salient tone), which did not entail prolonged decision processing and feedback anticipation (see Methods). The resulting response, termed ‘impulse response function’ (IRF) for simplicity, was more transient than those measured during the main experiment: the IRF returned back to the pre-stimulus baseline level after 3 s (Figure 3a, compare grey IRF with the blue line). Thus, the sustained elevations of pupil diameter observed beyond that time in the main experiment reflected top-down, cognitive modulations in pupil-linked arousal due to decision processing and reward anticipation (for the responses locked to the onset of the motion stimulus), or due to reward processing (for the feedback-locked responses). To quantify the amplitude of these cognitive modulations of the pupil response, we collapsed the pupil response across the time window 3-6 s from response (for pre-feedback interval) or from feedback tone (post-feedback interval; see gray shaded area in Figure 3a). For the cognitive modulations during the pre-feedback interval, we further extracted the mean pupil response values in the 500 ms before the feedback (gray shaded area in Figure 3b).

### Interacting effects of decision difficulty and accuracy on evoked pupil responses

The sustained pupil responses during both the intervals, pre- and post-feedback, scaled in line with the predictions from the Belief State Model, not the Stimulus State Model (compare Figure 3d-f with Figure 2b-d). First, pupil responses during both intervals were overall larger on error than correct trials (Figure 3b-c). The Stimulus State Model did not predict any difference between the two categories during the pre-feedback interval, because this model was only informed by external information (motion stimulus or feedback), not by noisy internal states. The larger pupil responses during errors in the pre-feedback interval were in line with previous results^3^, supporting the idea that arousal state between response and feedback reflects the observer’s decision uncertainty.

Second, the sustained pupil responses during both intervals exhibited a pattern of interactions between decision difficulty and accuracy as predicted by the Belief State Model but not the Stimulus State Model (compare Figure 3d to Figure 2d and Figure 2g). Hereby, the interaction was defined as (Easy Error - Easy Correct) - (Hard Error - Hard Correct). Specifically, the Belief State Model predicted a significant interaction of opposite sign for both intervals (Figure 2d, compare blue and purple dots). That same pattern was evident in the time course of the interaction term in the pupil response. During both intervals, the interaction terms were significant, with opposite signs: positive during the pre-feedback interval and negative during the post-feedback interval (Figure 3d, blue and purple bars). Consequently, the interaction terms were significantly different from one another throughout the entire part of the sustained pupil response (Figure 3d, black bar).

Finally, also the full pattern of sustained pupil responses for the Hard vs. Easy and Correct vs. Error conditions in both trial intervals (Figure 3e,f) resembled the pattern predicted by the Belief State Model (Figure 2b,c). In the sustained window during the post-feedback interval, there was a significant interaction between difficulty and accuracy (Figure 3f, *F*_(1,14)_ = 9.31, *p* = 0.009; Hard Error vs. Hard Correct, *p* = 0.001; Easy Error vs. Easy Correct, *p* = 0.174; Hard Error vs. Easy Error, *p* = 0.037; Hard Correct vs. Easy Correct, *p* = 0.031). The sustained window before feedback exhibited a trend towards an interaction (*F*_(1,14)_ = 4.12, *p* = 0.062). This effect became stronger during the pre-feedback interval: In the 0.5 s window just before (and locked to) the feedback delivery (grey window in Figure 3b), the interaction was significant (Figure 3e; *F*_(1,14)_ = 6.66, *p* = 0.022; post hoc comparisons: Hard Error vs. Hard Correct, *p* = 0.500; Easy Error vs. Easy Correct, *p* = 0.006; Hard Error vs. Easy Error, *p* = 0.037; Hard Correct vs. Easy Correct, *p* = 0.060). For all subsequent analyses, we focus on this interval 500 ms before feedback to probed into participants’ reward anticipation, referring to this time window as “pre-feedback interval”.

In sum, in this perceptual choice task, sustained pupil responses during both reward anticipation (pre-feedback) as well as after reward experience (post-feedback) were qualitatively in line with the predictions from a model of reward expectation and prediction errors model, in which the computation of these internal variables depended on internal belief states. The results from all main figures are only based on trials with long delay (> 7.5 s) intervals between choice and feedback, and feedback and subsequent trial, in order to minimize possible contamination of evoked pupil responses by responses to the next event (i.e., feedback or the next trial’s cue; see Methods). We found the same pattern of results when performing the analyses on trials (Supplementary Fig. S3).

### Control analysis for confounding effects of variations of RT and motion energy

In the current study, as in previous work using a similar perceptual choice task^3^, both RT and pre-feedback pupil dilation scaled with the decision uncertainty signal postulated by the Belief State Model. Indeed, RTs were significantly correlated to pre-feedback pupil responses in the pre-feedback window (−0.5-0 s) across all trials, *r*(13) = 0.12, *p* < 0.001, and within the following conditions: Hard Error, *r* = 0.11, *p* = 0.001; Hard Correct, *r* = 0.09, *p* < 0.001; Easy Correct, *r* = 0.16, *p* < 0.001, but not within the Easy Error condition, *r* = 0.07, *p* = 0.223.

While this association was expected under the assumption that RT and pupil dilation were driven by internal uncertainty signals^3^, the association also raised a possible confound. Arousal drives pupil dilation in a sustained manner throughout decision formation^25,34,35^. The peripheral pupil apparatus for pupil dilation (nerves and smooth muscles) has temporal low-pass characteristics. Consequently, trial-to-trial variations in decision time (the main source of RT variability) can cause trivial trial-to-trial variations in pupil dilation amplitudes, simply due to temporal accumulation of a sustained central input of constant amplitude but variable duration^25,34^. Then, pre-feedback pupil response amplitudes may have reflected RT-linked uncertainty, but without a corresponding scaling in the amplitudes of the neural input from central arousal systems. Note that this concern applied only to the pre-feedback pupil dilations, not the post-feedback dilations, which were normalized using the pre-feedback interval as baseline (see above). Another concern was that trial-by-trial fluctuations in motion energy, caused by the stochastically generated stimuli (see Methods) contributed to behavioral variability within the nominally Easy and Hard conditions.

Our results were not explained by either of those confounds (Figure 4). To control for both of them conjointly, we removed the influence of trial-to-trial variations in RT (via linear regression) from the pre-feedback pupil responses. And we used motion energy filtering^3,36^ to estimate each trial’s sensory evidence strength. We finally regressed the RT-corrected pupil time courses onto evidence strength (absolute motion energy), separately for the Error and Correct trials. The interaction term was defined as the difference in beta weights for the Error vs. Correct trial regressions. In this control analysis, the critical interaction effect was significant during both the pre-feedback and post-feedback time courses (*ps* < 0.05, cluster-based permutation test; Figure 4a). The interaction terms furthermore differed between intervals (*p* < 0.05, cluster-based permutation test; Figure 4a). When regressing mean RT-corrected pupil responses in the pre-feedback time window onto evidence strength, the critical interaction term (i.e. beta weights) within the pre-feedback window still reflected decision uncertainty (Figure 4b; *M* = 1.35, *STD* = 1.81, *p* = 0.001). In sum, while trial-to-trial variations in RT and motion energy explained some variance in the pupil responses, the key patterns of the pupil responses diagnostic of modulation by belief states were robust even when controlling for these parameters.

**Figure 4.**
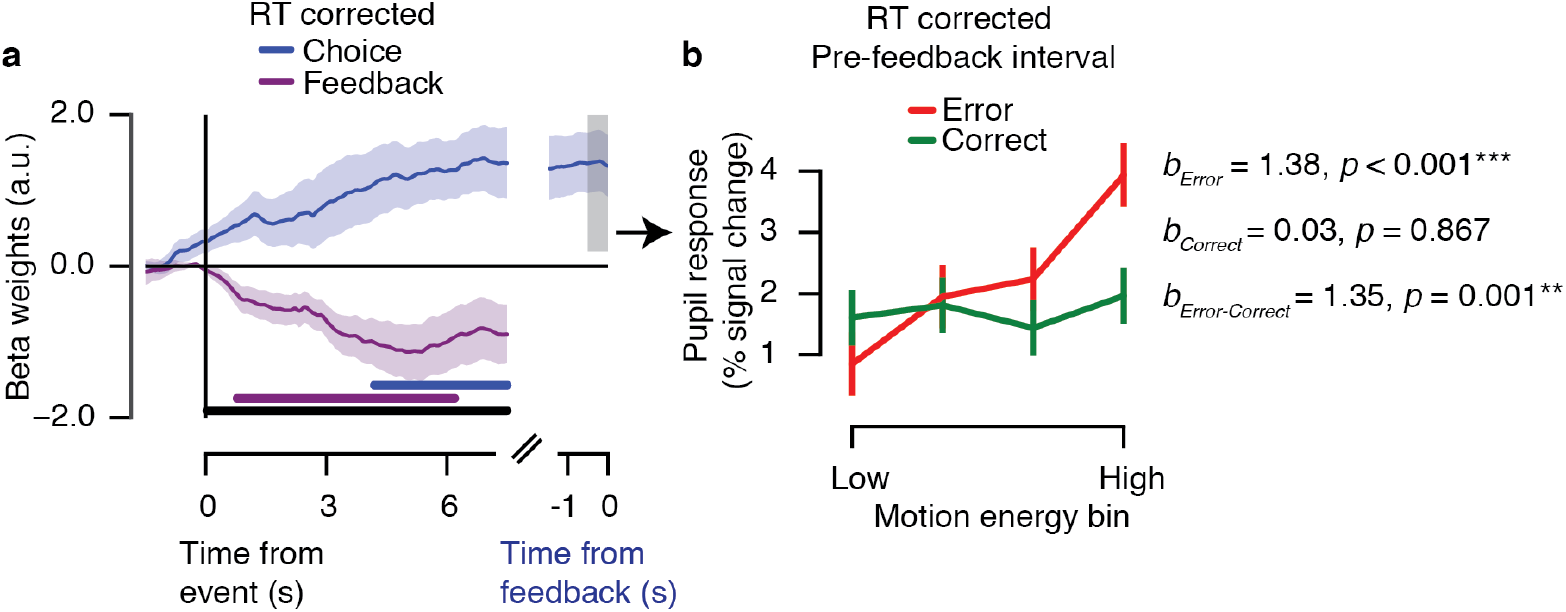
Pupil responses before feedback reflect observers’ belief state even when controlling for RT and motion energy fluctuations. **(a)** Time course of belief state scaling in the pupil, computed as trial-by-trial regression of RT-corrected pupil dilation onto motion energy strength. The interaction term (beta weights Error - beta weights Correct) is shown for choice-locked (blue, coinciding with onset of the choice) and feedback-locked (purple) responses. Horizontal bars indicate effect, *p* < 0.05 (cluster-based permutation test): blue bar indicates choice-locked response tested against 0; purple bar indicates feedback-locked response tested against 0; black bar indicates difference in interaction between both responses. **(b)** Mean response in pre-feedback time window (−0.5-0 s) as a function of difficulty and accuracy. Absolute motion energy was divided into four equally sized bins (per participant) for visualization. Error bars, standard error of the mean (*N* = 15). \**p* < 0.05, \*\**p* < 0.01, \*\*\**p* < 0.001.

### Relationship to Urai *et al*, 2017^3^

Our current results from the pre-feedback interval replicate our earlier finding^3^ that pupil responses in perceptual choice scale with decision uncertainty as postulated by the Belief State Model. This previous study focused on the pre-feedback responses and did not specifically assess the feedback-locked pupil responses (pupil measures were corrected with the same pre-trial baseline for the entire trial^3^). We here re-analyzed the post-feedback responses in the data from Urai et al. (2017) for comparison (see Supplementary Fig. S5). As in our current data, post-feedback responses were larger after incorrect than correct feedback (Supplementary Fig. S5a). However, the uncertainty-dependent scaling of post-feedback responses differed: rather than a negative interaction effect (Figure 1d), the interaction effect after feedback was positive (Supplementary Fig. S5b,c). One possible explanation for this difference the effect of reward-linked feedback: while participants in the current study were paid a compensation depending on their performance, feedback in the study by Urai et al. (2017) did not affect monetary reward. It is thus possible that the prospect of receiving performance-dependent monetary reward is required for the recruitment of pupil-linked arousal systems by uncertainty-dependent prediction errors. A number of further differences between these two studies complicate a direct comparison: the behavioral task (i.e. comparison of two intervals of motion strength vs. coarse direction motion direction discrimination), the short vs. long delay periods between events, and the two cohorts of participants. Despite these limitations, the difference in results between studies is potentially relevant and should be tested directly in follow-up work that eliminates the confounding factors listed above.

### Belief State Model predicts pupil responses quantitatively better than Stimulus State Model

The data presented thus far show that the pattern of pupil responses was qualitatively in line with the Belief State Model but not with the Stimulus State Model. To this end, we used predictions from model simulations based on the group data. However, individuals differ widely in terms of the internal noise, which dissociates between the models. We next tested whether the Belief State Model provides a quantitatively superior match to the measured pupil data than the Stimulus State Model when individual estimates of internal noise are used to generate model predictions. To this end, we simulated both models using individual estimates of internal noise (Supplementary Fig. S4a and Methods). This yielded model predictions for each individual for the Accuracy x Difficulty conditions, which were qualitatively in line with predictions based on the group, but with effects that varied in their magnitude between individuals depending on their estimated internal noise (Supplementary Fig. S4b).

We predicted that those individual patterns predicted by the Belief State Model should be more similar to the measured individual pupil responses than the individual patterns predicted by the Stimulus State Model. We tested this prediction by correlating predictions of both models with the corresponding pupil responses, separately for each individual. An example for a single subject is shown in Figure 5a, for both trial intervals. For both intervals, group-level correlations (Figure 5b) were significant (i.e. pupil responses similar) for the Belief State Model (*p* < 0.05), but not the Stimulus State Model (p > 0.05). Further, for the pre-feedback interval, the Belief State Model correlations were significantly stronger than the Stimulus State Model (*p* < 0.05). For the post-feedback interval, there was a trend towards a stronger correlation for the Belief State Model than the Stimulus State Model (*p* = 0.074).

**Figure 5.**
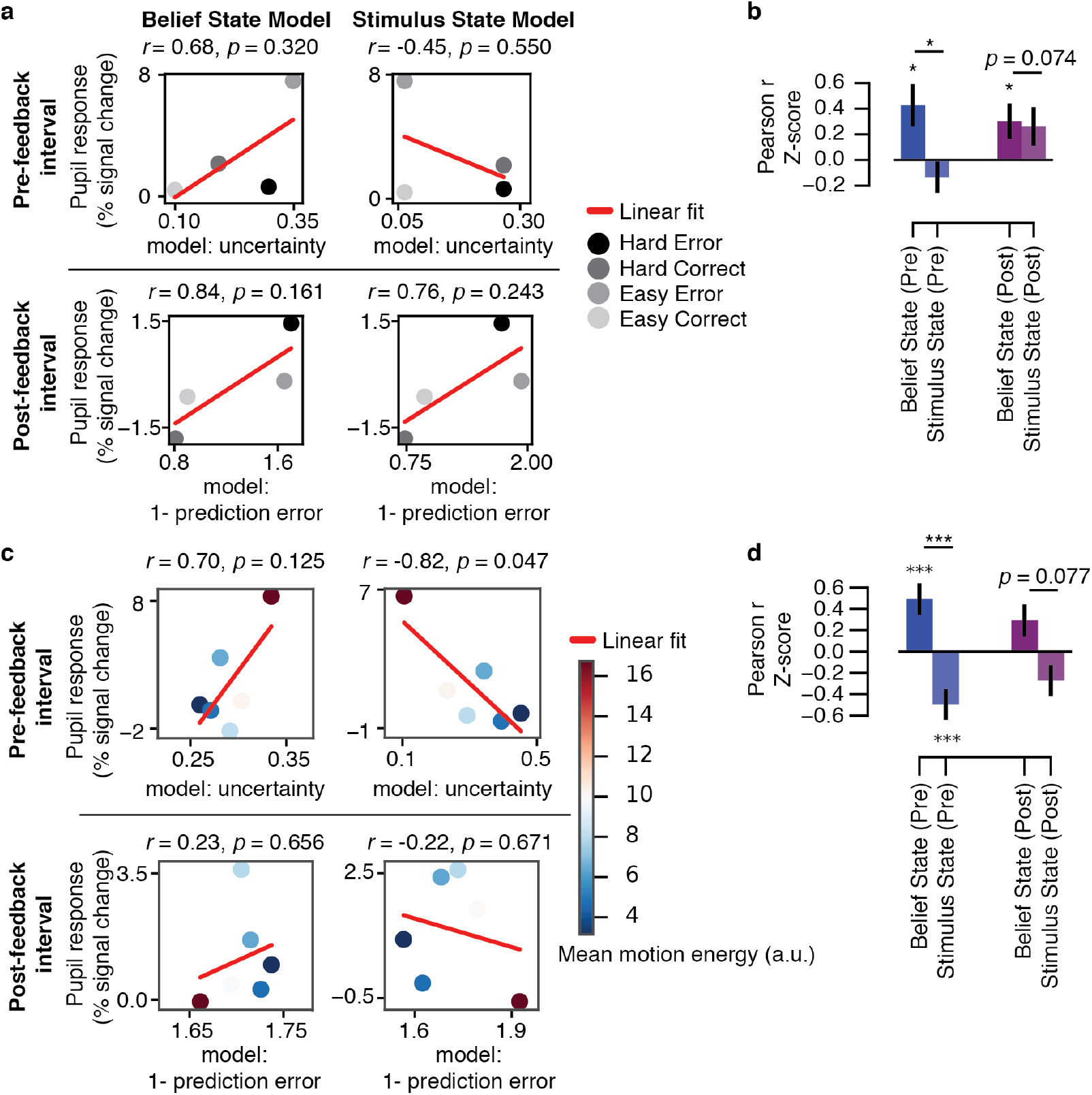
Model fits to pupil responses. **(a)** An example of the correlations (*r*) for a single subject. The four conditions of interest were defined by the Accuracy x Difficulty interaction. Easy and Hard conditions for the model parameters were averaged based on the coherence levels presented to each subject. **(b)** Group-level correlation coefficients (*r*) for the comparison of the model parameters and pupil responses, for the pre-feedback (Pre; −0.5-0 s) and post-feedback (Post; 3-6 s) intervals. **(c)** An example of the correlations for a single subject using model parameters simulated with motion energy (error trials only). Pupil responses were averaged within equal-sized bins based on the model parameter for each interval (6 bins). Evidence strength is represented by mean motion energy within each bin (color bar). **(d)** Group-level correlation coefficients (*r*) for the comparison of the model parameters (using motion energy) and pupil responses, for the pre-feedback (Pre; −0.5-0 s) and post-feedback (Post; 3-6 s) intervals (error trials only). Error bars, standard error of the mean (*N* = 15). \*\**p* < 0.01, \*\*\**p* < 0.001.

To perform a more fine-grained evaluation of the correspondence between model-predicted patterns and pupil responses, we used the motion energy information extracted from each trial (see previous section and Methods) rather than the categorical difficulty conditions (Easy, Hard) to generate individual model predictions. Because errors, not correct trials, qualitatively dissociate the predictions from Belief State and Stimulus State Models (compare Figure 2b,c with Figure 2e,f), we restricted this control analysis to error trials (Figure 5c,d). Again, predictions of both models were correlated to the corresponding pupil responses (6 bins of model parameters), separately for each individual. An example for a single subject is shown in Figure 5c.

For both intervals, correlations were positive (i.e. pupil responses similar) for the Belief State model predictions and negative (i.e. pupil responses dissimilar) for the Stimulus State model. Critically, the Belief State Model correlations were significantly larger than the Stimulus State Model in the pre-feedback interval (*p* < 0.001), again with a similar trend for the postfeedback interval (*p* = 0.074). The same held for a single-trial version of this correlation analysis, again focusing on error trials only (difference in correlation between models: *p* < 0.001 for pre-feedback; *p* < 0.080 for post-feedback).

## Discussion

It has long been known that the pupil dilates systematically during the performance of cognitive tasks^17–24^. The current study shows that task-evoked pupil dilation during a perceptual choice task indicates, at different phases of the trial, decision uncertainty and reward prediction error. Comparisons with qualitative model predictions showed that pupil responses during feedback anticipation and after reward feedback were modulated by decision-makers’ (noise-corrupted) internal belief states that also governed their choices. This insight is consistent with a reinforcement learning model (POMDP) that incorporates graded belief states in the computation of the prediction error signals^10,13^. In sum, the brain’s arousal system is systematically recruited in line with high-level computational variables.

A number of previous studies have related non-luminance mediated pupil responses to decision-making, uncertainty, and performance monitoring^14,15,34,37–41^, but our current results move beyond their findings in important ways. First, with the exception of Urai et al., (2017)^3^, previous studies linking uncertainty to pupil dynamics have used tasks in which uncertainty originated from the observer’s environment^14,15,37,39^. By contrast, in our task, decision uncertainty largely depended on the observers’ internal noise, which dissociated the two alternative models of the computational variables under study (decision uncertainty and reward prediction error, Figure 2). Second, our work went beyond the results from Urai et al., (2017)^3^ in showing that post-feedback pupil dilation reflects belief-modulated prediction error signals during perceptual decision-making in the context of monetary reward.

Previous work on central arousal systems and pupil-linked arousal dynamics has commonly used the dichotomy of (i) slow variations in baseline arousal state and (ii) rapid (so-called ‘phasic’) evoked responses^6,34,42,43^. Our current results indicate that this dichotomy is oversimplified, by only referring to the extreme points on a natural continuum of arousal dynamics during active behavior. Our results show that uncertainty around the time of decision formation as well as the subsequent reward experience both boost pupil-linked arousal levels in a sustained fashion: Pupils remained dilated for much longer than what would be expected from an arousal transient (Figure 3, compare all time courses with IRF). Even in our comparably slow experimental design, these sustained dilations lasted until long after the next experimental event. This implies that the sustained evoked arousal component we characterized here contributes significantly to trial-to-trial variations in baseline pupil diameter, which have commonly been treated as ‘spontaneous’ fluctuations.

Our insights are in line with theoretical accounts of the function of neuromodulatory brainstem systems implicated in the regulation of arousal^6,9^. Recent measurements in rodents, monkeys, and humans have shown that rapid pupil dilations reflect responses of neuromodulatory nuclei^25,28,44^. Neuromodulatory systems are interesting candidates for broadcasting uncertainty signals in the brain because of their potential of coordinating changes in global brain state^6,42^ and enabling synaptic plasticity in its target networks^45,46^. While pupil responses evoked by decision tasks or micro-stimulation have commonly been associated with the noradrenergic locus coeruleus^25,28,44,47,48^, these studies also found correlates in other brainstem systems^25,28,44^. In particular, task-evoked pupil responses during perceptual choice correlate with fMRI responses in dopaminergic nuclei, even after accounting for correlations with other brainstem nuclei (de Gee et al., (2017)^25^, their Figure 8H). Several other lines of evidence also point to an association between dopaminergic activity and nonluminance mediated pupil dilations. First, the locus coeruleus and dopaminergic midbrain nuclei are (directly and indirectly) interconnected^49–51^. Second, both receive top-down input from the same prefrontal cortical regions^49^, which might endow them with information about high-level computational variables such as belief states. Third, task-evoked fMRI responses of the locus coeruleus and substantia nigra are functionally coupled, even after accounting for correlations with other brainstem nuclei (de Gee et al., (2017)^25^, their Figure 8G). Fourth, both neuromodulatory systems are implicated in reward processing^48,50^. Fifth, rewards exhibit smaller effects on pupil dilation in individuals with Parkinson’s disease than in age-matched controls, a difference that can be modulated by dopaminergic agonists^52^. Future invasive studies should establish this putative link between pupil diameter and the dopamine system.

Recordings from midbrain dopamine neurons in monkey have also uncovered dynamics on multiple timescales^53,54^, in line with our current insights into pupil-linked uncertainty signaling. Further, the pattern of pupil dilations measured in the current study matched the functional characteristics of dopamine neurons remarkably closely (specifically, the pattern of the interaction between task difficulty and accuracy in pre- and post-feedback responses)^10^. However, the pupil responses followed the complement of the computational variables (i.e., 1-confidence and 1-prediction error) and the dopamine neurons identified by Lak et al., (2017)^10^. It is tempting to speculate that task-evoked pupil responses track, indirectly, the sign-inverted activity of such belief-state modulated dopaminergic system. Another alternative is that other brainstem systems driving pupil dilations^25,28,44^, exhibit the same belief-state modulated prediction error signals as dopamine neurons.

Our current work has some limitations, but also broader implications, which might inspire future work. First, provided that participants had learned the required (constant) decision boundary, the current task used did not require them to learn any environmental statistic. While prediction error signal such as the one studied here may be essential for perceptual learning^55,56^, the importance of the pupil-linked arousal signals for learning remains speculative in the context of our experiment. Future work should address their link to learning. In particular, while decision uncertainty can also be read out from behavioral markers such as RT^3,32,33^, no overt behavioral response is available to infer internal variables instantiated in response to feedback. Thus, our insight that the post-feedback pupil dilation reports a signal that is known to drive learning in the face of state uncertainty^13^ paves the way for future studies using this autonomous marker for tracking such learning signals in the brain.

Another important direction for future research is the relationship between pupil-linked uncertainty signals and the sense of confidence as reported by the observer^38^. The Belief State Model we used here makes predictions about a computational variable, statistical decision confidence^11^, while being agnostic about the mapping to the sense of confidence experienced or reported by the observer. Human confidence reports closely track statistical decision confidence in some experiments ^33^, but suffer from miscalibration in others, exhibiting over- or underconfidence^57^, insensitivity to the reliability of the evidence^58^, or biasing by affective value^59^.

In sum, we have established that internal belief states during perceptual decisionmaking, as inferred from a statistical model, are reflected in task-evoked pupil responses. This peripheral marker of central arousal can be of great use to behavioral and cognitive scientists interested in the dynamics of decision-making and reward processing in the face of uncertainty.

## Methods

An independent analysis of these data for the predictive power of pupil dilation locked to motor response, for perceptual sensitivity and decision criterion has been published previously^25^. The analyses presented in the current paper are conceptually and methodologically distinct, in that they focus on the relationship between Belief State Model predictions and pupil dilation, in particular locked to presentation of reward feedback.

### Participants

Fifteen healthy subjects with normal or corrected-to-normal vision participated in the study (6 women, aged 27 ± 4 years, range 23-37). The experiment was approved by the Ethical Committee of the Department of Psychology at the University of Amsterdam. All subjects gave written informed consent. All experiments were performed in accordance with the ethical guidelines and regulations. Two subjects were authors. Subjects were financially compensated with 10 Euros per hour in the behavioral lab and 15 Euros per hour for MRI scanning. In addition to this standard compensation, subjects earned money based on their task performance: 0-10 Euros linearly spaced from 50-100% accuracy per experimental session (i.e. 50% correct = 0 Euros, 75% = 5 Euros, 100% = 10 Euros). At the end of each block of trials, subjects were informed about their average performance accuracy and corresponding monetary award. Earnings were averaged across all blocks at the end of each session.

### Behavioral task and procedure

Subjects performed a two-alternative forced choice (2AFC) motion discrimination task while pupil dilation was measured (Figure 1). Motion coherence varied so that observers performed at 70% correct in 2/3 of trials (‘hard’) and at 85% correct in 1/3 of trials (‘easy’). After a variable delay (3.5-11.5 s) following the choice on each trial, we presented feedback that was coupled to a monetary reward (see ‘Participants’).

Each subject participated in one training session and four main experimental sessions (in the MRI scanner). During the training session, subjects’ individual threshold coherence levels were determined using a psychometric function fit with 7 levels, 100 trials per level, 0-80% coherence. The training session took 1.5 hours and each experimental session lasted 2 hours. During the experimental sessions, stimuli were presented on a 31.55” MRI compatible LCD display with a spatial resolution of 1920 × 1080 pixels and a refresh rate of 120 Hz.

The individual coherence levels were validated at the beginning of each experimental session in practice blocks (during anatomical scans) by checking that the subject’s average accuracy across a block corresponded to 75% correct. If subjects’ average accuracy of a block exceeded 75%, the difficulty of the task was increased in the following block by slightly decreasing the motion coherence based on individual performance thresholds (in steps of 1% in accuracy, equally for both Hard and Easy conditions). During experimental blocks, greater motion coherence (i.e. stronger evidence strength) resulted in higher accuracy as well as faster responses. Subjects’ accuracy was higher on Easy trials (*M* = 88.06% correct, *SD* = 4.26) compared to Hard trials (*M* = 71.15% correct, *SD* = 3.64), *p* < 0.001. Subjects were faster to respond on Easy trials (*M* = 1.13 s, *SD* = 0.13) compared to Hard trials (*M* = 1.22 s, *SD* = 0.14), *p* < 0.001.

Task instructions were to indicate the direction of coherent dot motion (upward or downward) with the corresponding button press and to continuously maintain fixation in a central region during each task block. Subjects were furthermore instructed to withhold responses until the offset of the coherent motion stimulus (indicated by a visual cue). The mapping between perceptual choice and button press (e.g., up/down to right/left hand button press) was reversed within subjects after the second session (out of four) and was counterbalanced between subjects. Subjects used the index fingers of both hands to respond.

Each trial consisted of five phases during which random motion (0% coherence) was presented, with the exception of the stimulus interval: (i) the pupil baseline period (0.5-7 s); (ii) the stimulus interval consisting of random and coherent motion for a fixed duration of 0.75 s; (iii) the response window (maximum duration was 2.25 s); (iv) the delay period preceding feedback (3.5-11.5 s, uniformly distributed across 5 levels); (v) the feedback and the inter-trial interval (ITI; 3.5-11.5 s, uniformly distributed across 5 levels). Stimulus onset coincided with a visual and auditory cue. The auditory cue was presented for 0.25 s (white noise or pure tone at 880 Hz, 50-50% of trials, randomly intermixed). The visual cue was a change in the region of fixation from an open to a closed rectangle. The return of the fixation region to an open rectangle indicated to subjects to give their response (the surface areas in pixels of the open and closed rectangles were held equal in order to assure no change in overall luminance). Feedback was presented visually (green/red for correct/error) for 50 frames (0.42 s at 120 Hz). If subjects did not respond or were too fast/slow in responding, a yellow rectangle was presented as feedback on that trial.

Each block of the task began and ended with a 12-s baseline period, consisting of a fixation region (no dots). Each block of the task had 25 trials and lasted approximately 8 minutes. Subjects performed between 23 and 24 blocks yielding a total of 575–600 trials per subject. One subject performed a total of 18 blocks (distributed over three sessions), yielding a total of 425 trials. Data from one session of two subjects (12 blocks in total) and 2 blocks of a third subject were excluded from the analyses because of poor eye-tracker data quality or technical error.

### Visual stimuli

Dot motion stimuli were presented within a central annulus that was not visible to the subjects (grey background, outer diameter 16.8°, inner diameter of 2.4°). The fixation region was in the center of the annulus and consisted of a black rectangle (0.45° length). Signal dots moved at 7.5°/s in one of two directions (90° or 270°). Noise dots were randomly assigned (uniformly distributed) to locations within the annulus on each frame, preventing them from being trackable. Each frame consisted of 524 white dots (0.15° in diameter) within one visual hemifield (left or right; The hemifield remained constant during a block of trials and was counterbalanced between blocks. This manipulation was specific for the MRI experiment; the two hemifields were averaged in the current analysis). The proportion of ‘signal’ as compared with ‘noise’ dots defined motion coherence levels. Signal dots were randomly selected on each frame, lasted 10 frames, and were thereafter re-plotted in random locations (reappearing on the opposite side when their motion extended outside of the annulus). To prevent tracking of individual dots, independent motion sequences (n = 3) were interleaved^60^.

### Eye-tracking data acquisition and preprocessing

Pupil diameter was measured using an EyeLink 1000 Long Range Mount (SR Research, Osgoode, Ontario, Canada). Either the left or right pupil was tracked (via the mirror attached to the head coil) at 1000 Hz sample rate with an average spatial resolution of 15 to 30 min arc. The MRI681 compatible (non-ferromagnetic) eye tracker was placed outside the scanner bore. Eye position was calibrated once at the start of each scanning session.

Eye blinks and saccades were detected using the manufacturer’s standard algorithms (default settings). Further preprocessing steps were carried out using custom-made Python software, which consisted of (i) linear interpolation around blinks (time window from 0.1 s before until 0.1 s after each blink), (ii) band-pass filtering (third-order Butterworth, passband: 0.01–6 Hz), (iii) removing responses to blink and saccade events using multiple linear regression (responses estimated by deconvolution)^61^, and (iv) converting to percent signal change with respect to the mean of the pupil time series per block of trials.

### Quantifying pre- and post-feedback pupil responses

Pupil dilation is affected by a range of non-cognitive factors^51^, whose impact needs to be eliminated before inferring the relation between central arousal and computational variables of interest. We excluded the impact of a number of non-cognitive factors on the pupil responses: (i) blinks and eye movements, which were eliminated from the analysis (see above); (ii), luminance, which was held constant throughout the trial, with the exception of the visual feedback signals, which we controlled for in a separate control experiment: Supp. Fig. S2); (iii) motor responses^62^; and (iv) trial-by-trial variations in decision time that may confound pupil response amplitudes^25,34^ due to the temporal accumulation properties of the peripheral pupil apparatus^63,64^. With the aim of excluding effects related to above mentioned points (iii) and (iv), we investigated pupil responses locked to the choice reported by the observer. Additionally, only trials with the three longest delay intervals between events (7.5, 9.5 and 11.5 s; 3/5 of all trials) were used in the main analysis of pupil responses. Specifically, for the pre-feedback interval, the delay period was between the choice and feedback. For the post-feedback interval, the delay period was the inter-trial interval. Finally, we performed a control analysis in which RTs are removed from pupil responses via linear regression (see Figure 4).

For each trial of the motion discrimination task, two events of interest were inspected: (a) pupil responses locked to the observers’ reported choice (button press) and (b) pupil responses locked to the onset of the feedback. On each trial, the mean baseline pupil diameter (the preceding 0.5 s) with respect to the motion stimulus onset and feedback was subtracted from the evoked response for each event of interest on each trial. We extracted the mean pupil responses within the sustained time window (3-6 s), defined by the period during which the independently measured pupil IRF returned to baseline (at the group level, Figure 3a). The uncertainty signal was expected to be largest in the time window just preceding feedback based on Urai et al. (2017)^3^, reflecting the fact that the ‘reward anticipation’ state is highest the longer the observer waits for feedback. Therefore, we additionally analyzed pre-feedback pupil responses in the 0.5 s preceding feedback.

### Model predictions

In signal detection theory, on each trial a decision variable (*dv_i_*) was drawn from a normal distribution *N*(*μ, σ*), where *μ* was the sensory evidence on the current trial and *σ* was the level of internal noise. In our case, we took *μ* to range from −0.5 to 0.5, corresponding to the extremes of the motion coherence presented in the main experiment (where 0 = 100% random motion and 1 = 100% coherent motion). The internal noise, *σ*, was estimated by fitting a probit psychometric function onto the combined data across all subjects (slope *β* = 7.5). The standard deviation, *σ*, of the *dv* distribution is 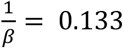. The decision bound, *c*, was set to 0, indicating no choice bias for any observer.

For each level of evidence strength, *μ* = [-0.5, 0.5] in steps of 0.01, we simulated a normal distribution of *dv* with *σ* = 0.133 with 10,000 trials. The choice on each trial corresponded to the sign of *dv_i_*. A choice was correct when the sign of *dv_i_* was equal to the sign of *μ_i_*. Errors occurred due to the presence of noise in the *dv*, which governed choice in both of the two models discussed as follows.

We simulated two models, Belief State Model and Stimulus State Model, which differed only in the input into the function used to compute confidence: whether the confidence is a function of *dv¿* or μ¿. Confidence was defined as

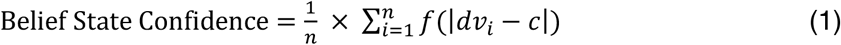

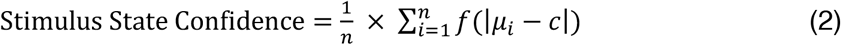

where n was the number of trials per condition, for which the predictions were generated (see below), *f* was the cumulative distribution function of the normal distribution, transforming the distance |*dv* – *c*| or |*μ* – *c*| into the probability of a correct response, for the Belief State or Stimulus State Model, respectively

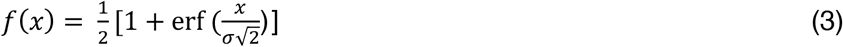

Because we applied equations 1 and 2 separately to each combination of Difficulty (i.e. coherence level) and Accuracy (Error and Correct) conditions, *n* depended on the variable number of trials obtained in each condition (with the smallest *n* for Easy Error) in our simulations.

Decision uncertainty was the complement of confidence

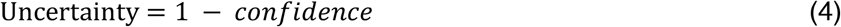

And the single-prediction error was defined as

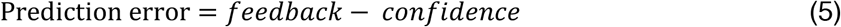

where *feedback* was 0 or 1.

Pre-feedback pupil responses have previously been found to reflect decision uncertainty^3^; we therefore expected the post-feedback pupil responses to similarly follow the complement of the prediction error (i.e. 1-*prediction error*). For each trial, we computed the binary choice, the level of decision uncertainty, the accuracy of the choice and the prediction error. For plotting, we collapsed the coherence levels across the signs of *μ*, as these are symmetric for the up and down motion directions.

Custom Python code used to generate the model predictions can be found here: https://github.com/colizoli/pupil_belief_states.

### Motion energy

To extract estimates of fluctuating sensory evidence, we applied motion energy filtering to the single-trial dot motion stimuli (using the filters described in Urai and Wimmer, 2016^36^). Summing the 3D motion energy values over space and time gave us a single-trial estimate of the external sensory evidence presented to the subject (positive for upwards, negative for downwards motion). We used the absolute value of this signed motion energy signal as our continuous measure of sensory evidence strength in statistical analyses. for visualization (Figure X), we divided this absolute motion energy metric into 4 equally-sized bins within every observer.

### Statistical analysis

Behavioral variables and pupil responses were averaged for each condition of interest per subject (*N* = 15). Statistical analysis of mean differences in pupil dilation of evoked responses was done using cluster-based permutation methods^65^. The average responses in the sustained time window were evaluated using a two-way ANOVA with factors: difficulty (2 levels: Hard vs. Easy) and accuracy (2 levels: Correct vs. Error). All post-hoc and two-way comparisons were based on non-parametric permutation tests (two-tailed).

### Control experiment 1: Individual pupil impulse response functions

In order to define a sustained component of pupil responses evoked by the events of interest during the main experiment, we independently measured subjects’ pupil responses evoked by simply pushing a button upon hearing a salient cue. This enabled a principled definition of the time window of interest in which to average pupil responses based on independent data. Subjects performed one block of the pupil impulse response task at the start of each experimental session (while anatomical scans were being acquired). Pupil responses following an auditory cue were measured for each subject^63^. Pupils were tracked while subjects maintained fixation at a central region consisting of a black open rectangle (0.45° length) against a grey screen. No visual stimuli changed, ensuring constant illumination within a block. An auditory white noise stimulus (0.25 s) was presented at random intervals between 2 and 6 s (drawn from a uniform distribution). Participants were instructed to press a button with their right index finger as fast as possible after each auditory stimulus. One block consisted of 25 trials and lasted 2 min. Two subjects performed three blocks, yielding a total of 75-100 trials per subject. Trials without a response were excluded from the analysis. Each subject’s impulse response function (IRF) was estimated using deconvolution (with respect to the auditory cue) in order to remove effects of overlapping events due to the short delay interval between subsequent trials^61^.

### Control experiment 2: Pupil responses during passive viewing of feedback signals

Pupil responses evoked by the green and red fixation regions used in the main experiment were measured in a separate control experiment (see Supplementary Fig. S2; *N* = 15, 5 women, aged 28.5 ± 4 years, range 23-34). Three subjects were authors, two of which participated in the main 2AFC task. No other subjects from this control experiment participated in the main 2AFC task. Pupils were tracked while subjects maintained fixation at a central region of the screen. Stimuli were identical to the main 2AFC task; dot motion consisted of only random motion (0% coherence). A trial consisted of two phases: (i) the baseline period preceding the onset of a color change (1-3 s, uniform distribution), and (ii) passive viewing of the stimuli used for feedback in the main experiment: during which the fixation region changed to either red or green (50-50% of trials, randomized) for 50 frames (0.42 s at 120 Hz). This was followed by an ITI (3-6 s, uniformly distributed). Participants were instructed that they did not need to respond, only to maintain fixation. A block consisted of 25 trials and lasted 3 min. Subjects performed eight blocks of this task in the behavioral lab, yielding 200 trials per subject.

## Acknowledgments

This research was supported by the European Union Seventh Framework Programme (FP7/2007-2013) under grant agreement no. 604102 (Human Brain Project) (to T.H.D.), the German Research Foundation (DFG): DO 1240/3-1, DO 1240/2-1, and SFB 936/A7 (to T.H.D.) and the German Academic Exchange Service (DAAD) (to A.E.U.).

## Author contributions

O.C. contributed Conceptualization, Formal analysis, Investigation, Writing - original draft, Writing - review and editing; J.W.G. contributed Conceptualization, Formal analysis, Investigation, Writing - original draft, Writing - review and editing; A.E.U. contributed Formal analysis, Writing - original draft, Writing - review and editing; T.H.D. contributed Conceptualization, Resources, Supervision, Writing - original draft, Writing - review and editing;

## Data availability

The pupil data and model prediction code are publicly available here: https://github.com/colizoli/pupil_belief_states.

## Materials & Correspondence

(T.H.D.)

---

**Supplementary Figure S1.**
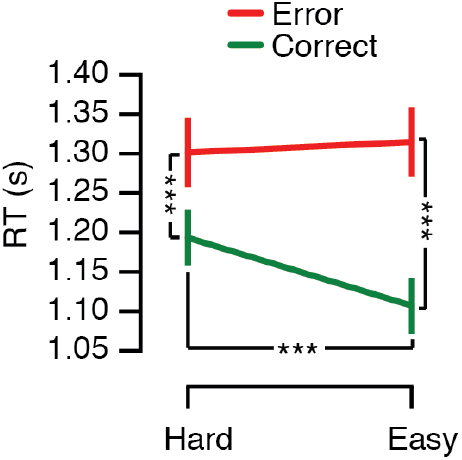
RT scales with decision uncertainty. Mean reaction times (RT) as a function of task difficulty and accuracy. Task difficulty and accuracy interacted. Error bars represent the standard error of the mean (*N* = 15). \*\*\**p* < 0.001

**Supplementary Figure S2.**
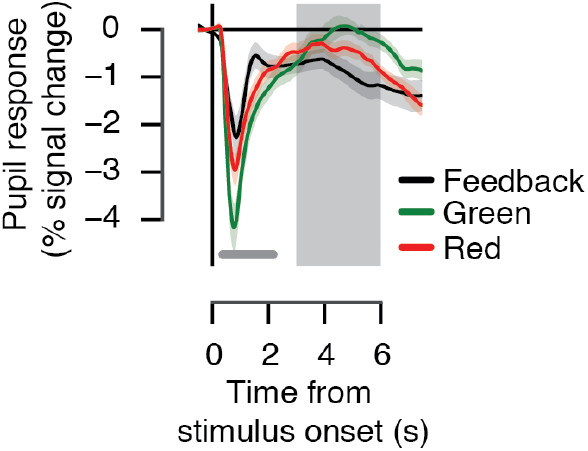
Pupil responses during passive viewing of feedback signals. In a control experiment (*N* = 15, 5 women, aged 28.5±4 years, range 23-34), we investigated the time course of potential differences in pupil responses evoked by red as compared with green light, regardless of whether these colors correspond to reward feedback during the perceptual choice task. Three subjects were authors, two of which participated in the main experiment. Stimuli were identical to the main 2AFC task; dot motion consisted of only random motion (0% coherence). A trial consisted of a baseline period preceding the onset of a color change (1-3 s, uniformly distributed) the red or green dot at fixation, and ITI (3-6 s, uniformly distributed). Participants passively viewed the stimuli while maintaining fixation. Pupil responses were averaged for each condition of interest per subject (*N* = 15, 200 trials per subject). The light-mediated pupil constrictions evoked by visual feedback cues during main task (grey) and in passive viewing control experiment (red, green). The grey bar indicates a difference between red- and green-evoked responses, *p* < 0.05 (cluster-based permutation test, see main text). Grey shaded area, ‘sustained’ time window during which pupil dilation was averaged, defined by the period during which the pupil impulse response function returned to baseline and the shortest delay between events (3-6 s). The results show that (i) green and red light both evoked pupil constrictions, and (ii) green light produced slightly larger pupil constriction than red light, in an early time window (0.25-2.25 s). The difference in correct vs. error trials in pupil constriction after feedback during the main experiment continues after this early time window (see Figure 3c). Furthermore, any differences obtained *within* error and correct conditions after feedback during the main experiment cannot be explained by differences between the color-evoked responses, as the stimulus color was the same between these comparisons.

**Supplementary Figure S3.**
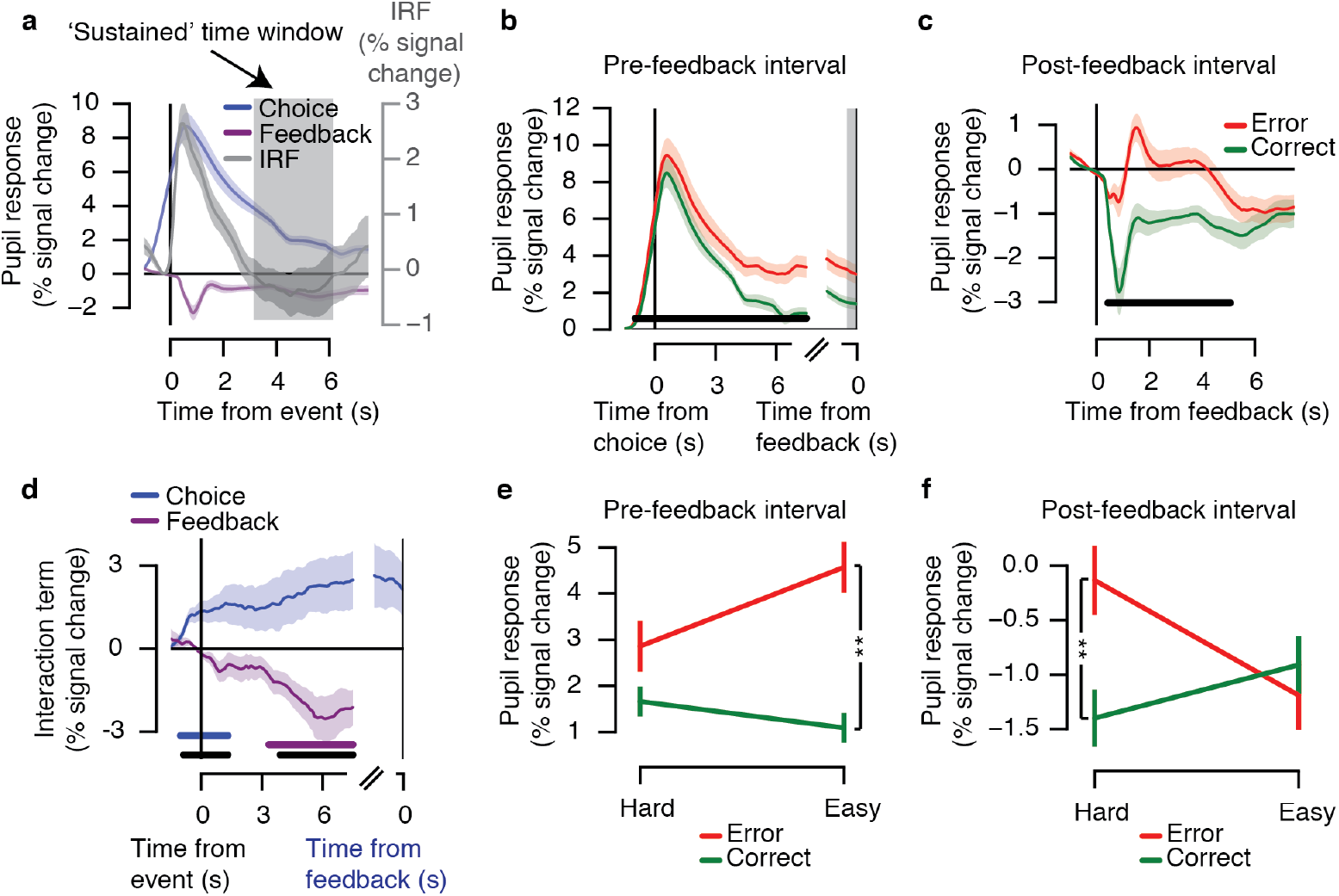
Replication of Figure 3 with all trials. The same pattern of pupil responses was obtained when all trials, including those in which the interval between response and feedback was less than 3 s, were included in the analysis **(a-f)**. For the pupil responses in the −0.5 s window preceding feedback **(e)**, a significant interaction between difficulty and accuracy was obtained in this later time window (*F*_(1,14)_ = 4.95, *p* = 0.043; post hoc comparisons: Hard Error vs. Hard Correct, *p* = 0.072; Easy Error vs. Easy Correct, *p* = 0.004; Hard Error vs. Easy Error, *p* = 0.074; Hard Correct vs. Easy Correct, *p* = 0.061). During the post-feedback interval, a significant interaction between difficulty and accuracy was obtained (f), *F*_(1,14)_ = 7.89, *p* = 0.014; Hard Error vs. Hard Correct, *p* = 0.001; Easy Error vs. Easy Correct, *p* = 0.572; Hard Error vs. Easy Error, *p* = 0.066; Hard Correct vs. Easy Correct, *p* = 0.118).

**Supplementary Figure S4.**
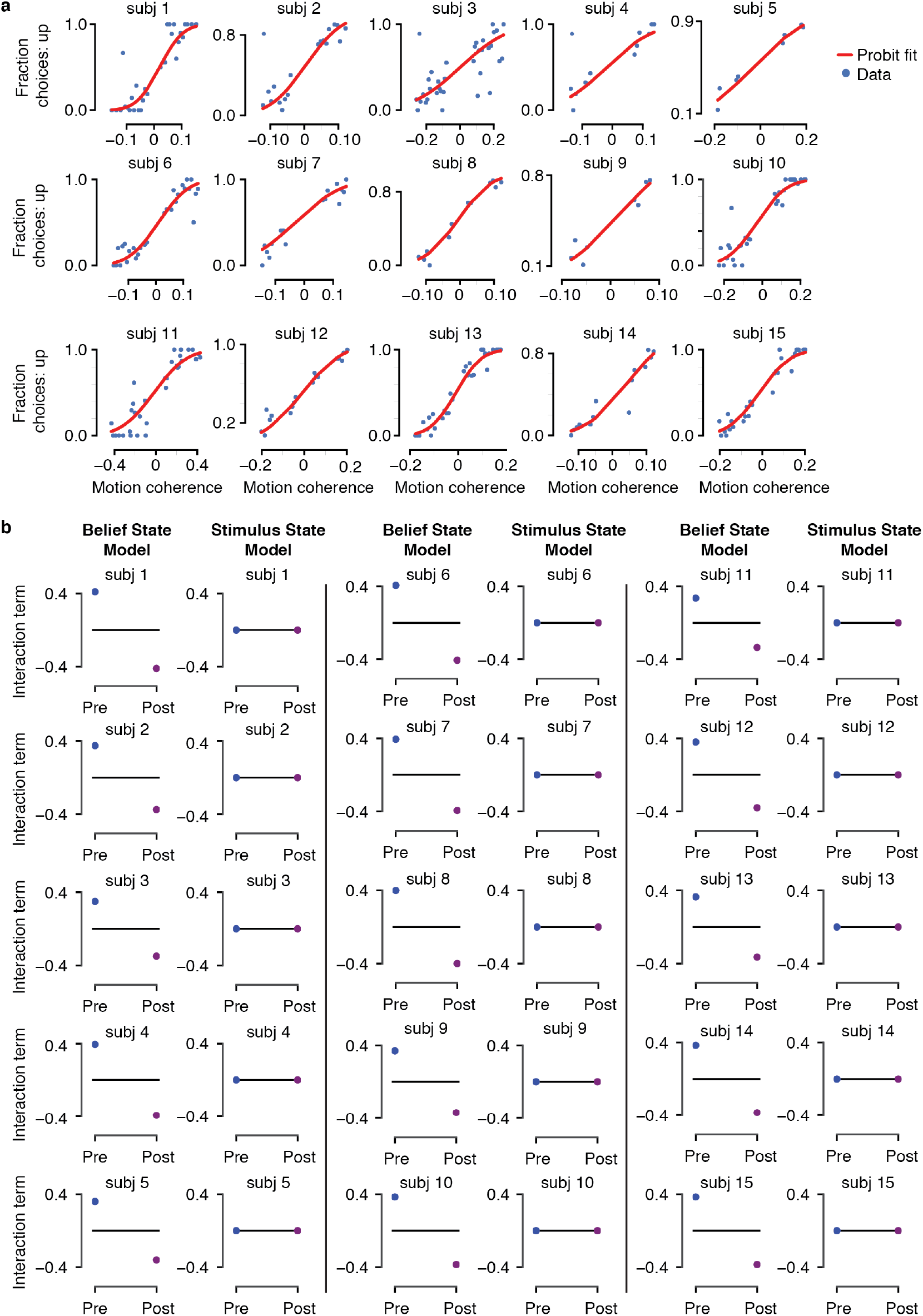
Individual psychometric functions and model predictions. Based on the individual estimates of internal noise in the data (i.e. sigma) **(a)**, subject-specific model predictions were generated for the Belief State and Stimulus State models **(b)**. Predictions for the interaction term defined as (Easy Error - Easy Correct) - (Hard Error - Hard Correct) based on subject-specific motion coherence levels are shown.

**Supplementary Figure S5.**
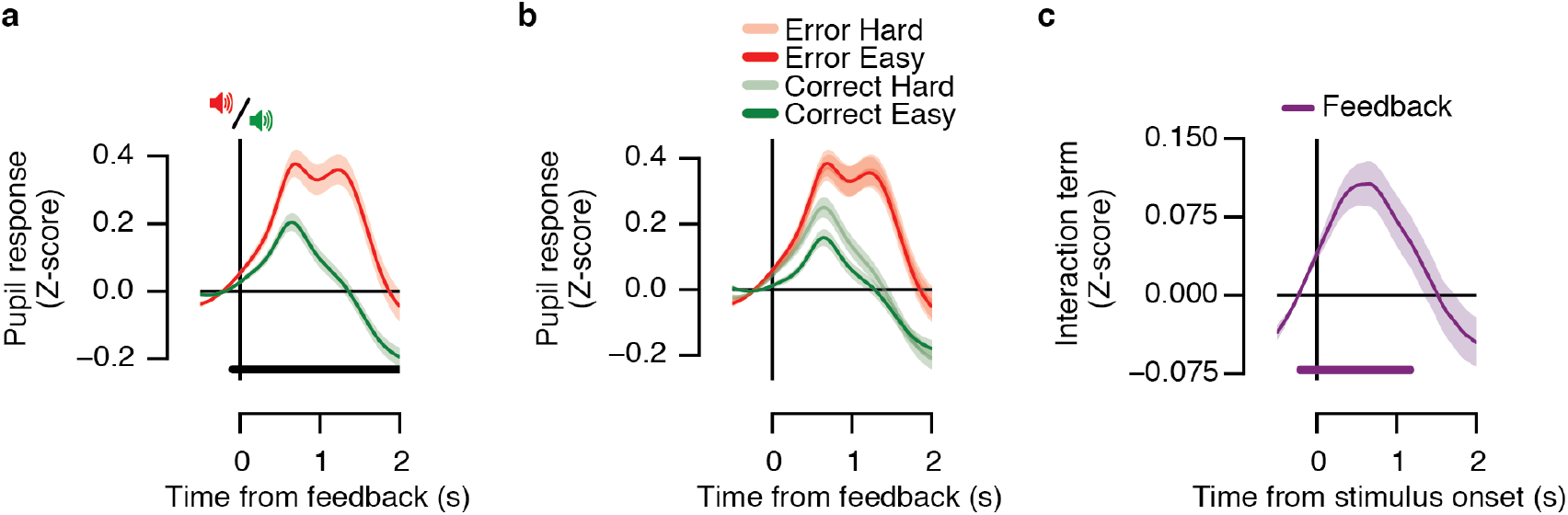
Feedback-locked responses from Urai et al. (2017). Reanalysis of the data from our previously published study (available at https://doi.org/10.6084/m9.figshare.4300043). This study used a similar visual perceptual choice task, however with a number of important differences specified in the following: The study used a two-interval forced choice motion coherence discrimination task; multiple levels of task difficulty were intermixed, here sorted into two categories (median split) yielding Hard and Easy conditions for comparison with the present data; delay intervals between decision and feedback, and the inter-trial-intervals were shorter than in the current study; feedback (Correct or Error) was presented by two different tones; feedback was not linked to any reward (participants’ financial remuneration was not contingent on performance). **(a)** Evoked pupil responses for Correct (green) and Error (red) trials locked to trial-wise (auditory) feedback. The black bar indicates correct vs. error effect, *p* < 0.05 (cluster-based permutation test). Because feedback was not presented visually, there was no post-feedback pupil constriction, but dilation for all trial types. Error feedback elicited stronger dilations than correct feedback, as in the current data (compare with Figure 3c). **(b)** Pupil responses as a function of task difficulty and accuracy locked to feedback. The scaling with evidence strength was similar to pre-feedback decision uncertainty, but not to post-feedback prediction error, with smaller dilations for Correct Easy than Correct Hard responses (compare to Figure 2c). **(c)** The interaction term for task difficulty with two levels (Easy Error - Easy Correct) - (Hard Error - Hard Correct) for feedback-locked responses. The purple bar indicates feedback-locked response tested against 0, *p* < 0.05 (cluster-based permutation test). For all feedback-locked responses, the mean pupil diameter across the pre-feedback interval from −0.5 s to 0 s was subtracted from the response time courses at the single-trial level. Each condition of interest was averaged across subjects (*N* = 27).

